# Seasonally variable thermal performance curves prevent adverse effects of heatwaves

**DOI:** 10.1101/2023.05.09.540050

**Authors:** Matthew C. Sasaki, Michael Finiguerra, Hans G. Dam

## Abstract

The increasing frequency and intensity of heatwaves may represent a significant challenge for predicting vulnerability of populations in a warming ocean. The direct impacts of heatwaves on populations depend on the relative position of environmental temperatures to the thermal performance curve optima. If thermal performance curves are static, the effects of heatwaves may therefore change seasonally over the annual temperature cycle. However, these seasonal changes in the effects of heatwaves may be dampened by corresponding variation in thermal performance curves which, in organisms with relatively short generation times, may be driven by phenotypic plasticity as well as genetic differentiation. Here we investigate the effects of seasonal timing and duration on the impacts of heatwaves in the ecologically important copepod congeners *Acartia tonsa* and *Acartia hudsonica*, and test the hypotheses that 1) seasonal variation in thermal performance curves will reduce overall population vulnerability to heatwaves, and 2) that seasonal variation in TPCs will prevent negative transgenerational effects of heatwave. We characterized seasonal variation in thermal performance curves for several fitness-related traits. These experiments uncovered strong seasonal variation in the thermal performance curves of *Acartia tonsa*, and indicate that this variation buffers against negative effects of simulated heatwaves. We also quantified both direct and trans-generational effects of different duration heatwaves on copepods collected at various times throughout the season using simulated heatwave experiments. There was no consistent pattern in the transgenerational effects of parental exposure to heatwaves, which may indicate that seasonal variation in thermal performance curves reduces the effects of parental stress on offspring performance. Our results show that seasonal variation in thermal performance curves will likely play an important role in limiting the adverse effects of heatwaves on populations.

## Introduction

Heatwaves are increasing in frequency and intensity in aquatic ecosystems (Frölicher & Laufkötter, 2018; Oliver *et al*., 2018; Tassone *et al*., 2022). These periods of anomalously high temperatures present severe challenges to marine biota (Smale *et al*., 2019). The effects of these events on communities will be strongly dependent on the relative vulnerabilities of the different component taxa to increased temperatures. Understandably, much attention has been focused on macro-organisms like coral, kelp, and fish, on which heatwaves have had significant, highly visible effects (Smith *et al*., 2023). Planktonic taxa have received less attention, despite their importance in aquatic communities and unique life histories. Shaped by resulting changes in metabolism, reproduction, competitive ability, and survival, however, increased temperature during a heatwave may by expected to have have profound effects on planktonic community dynamics and the distribution of organisms (McKinstry *et al*., 2022; Evans *et al*., 2020; Maazouzi *et al*., 2008).

Characterizing the thermal sensitivity of planktonic taxa is crucial for predicting the effects of marine heatwaves on food webs and ecosystem functioning. Thermal performance curves provide key insights into the direct effects of increased temperature on organisms and population dynamics (Deutsch *et al*., 2022; Arroyo *et al*., 2022; Hochachka & Somero, 2002; Angilletta, 2009). However, in taxa with short generation times, seasonal acclimatization and genetic differentiation can rapidly change thermal performance curves to track environmental temperatures, modifying the effects of heatwaves (Sasaki & Dam, 2020; Rudman *et al*., 2022). Further, the effects of heatwaves on populations may also be propagated across generations by mechanisms like trans-generational plasticity and maternal effects, which can have important effects on offspring performance (Truong *et al*., 2022; Dinh *et al*., 2021; Minuti *et al*., 2022). These fine-scale temporal dynamics need to be accounted for when predicting the effects of heatwaves on populations of planktonic and other short-lived taxa.

Vulnerability to adverse effects of heatwaves is often assumed to be largest during the warmest seasons when environmental temperatures are nearest to organismal thermal optima. This can be modified, however, by seasonal variation in thermal performance (Tran & Johansen, 2023). In planktonic taxa, this seasonal variation may be underlain by acclimatization within generations, transgenerational effects across generations, and seasonal genetic differentiation (Sasaki & Dam, 2020). Regardless of mechanism, this variation may play an important role in determining the effects of heatwaves on a population (Figure 1). Much of the past work on the effects of heatwaves assumes performance curves are fixed (i.e. - there is no seasonal variation in TPCs). In this case, the effect of heatwaves would be predicted to change over the course of the season the relative position of environmental temperatures to TPC optimum temperature and lethal thermal limits varies. Alternatively, variation in TPCs over seasonal timescales may act as a buffer against negative effects of heatwaves if optimum temperatures maintain a fixed position above environmental temperatures, limiting exposure to temperatures in excess of thermal optima. Differences in the magnitude of seasonal variation in TPCs between taxa may therefore play a key role in shaping patterns in relative vulnerability of community members to heatwaves.

**Figure 1:**
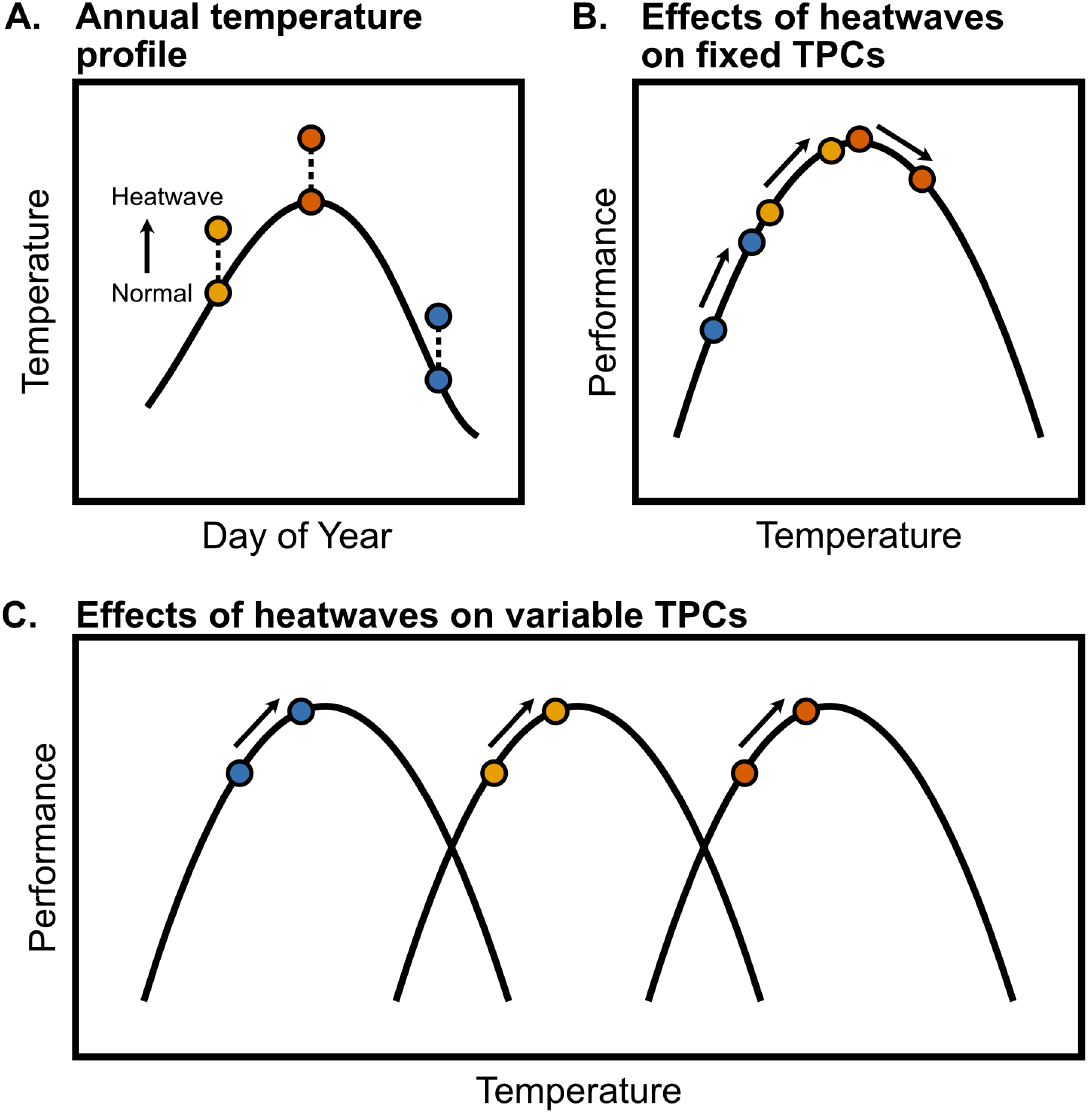
Schematic illustrating the effects of heatwaves on performance. A) Heatwaves may occur at various times throughout the year, resulting in acute increases in temperature against a seasonally-variable baseline temperature (shown here for an hypothetical location in the Northern Hemisphere). B) Static thermal performance curves result in seasonally-variable effects of heatwaves on performance. Heatwaves that occur when temperatures are low relative to the optimum would be expected to increase performance (a beneficial effect). Heatwaves that occur when temperatures are near or past the thermal optimum may decrease performance (a detrimental effect). C) Variable thermal performance curves reduce seasonal variation in the effects of heatwaves on performance, when thermal optima are positively correlated with environmental temperatures. An increase in temperature would then be expected to increase performance, regardless of the seasonal timing of the heatwave.

Given their short generations times, abundance, and ecological importance in marine systems (Dam, 2013), copepods are an ideal model system to examine the interplay between seasonal variation in thermal performance curves and the direct and indirect effects of heatwaves. Here we characterize how thermal performance curves for key fitness related traits (egg and offspring production, hatching success, and survivorship) of the ecologically important copepod species *Acartia hudsonica* and *A. tonsa* change over the season of occurrence, and how this variation shapes vulnerability to heatwaves. We also quantified both direct and trans-generational effects of heatwaves on one of the species (*A. tonsa*) in the laboratory using a series of simulated heatwave experiments. We test two hypotheses: 1) there is seasonal variation in the thermal performance curves of *A. hudsonica* and *A. tonsa*; and 2) that seasonal variation in TPCs will prevent negative transgenerational effects of heatwaves.

## Methods

### Generating Field TPCs

Copepods were collected from eastern Long Island Sound, USA (41.31, -72.07) several times throughout the year using surface tows of a 200 um mesh plankton net (Table 1). Mature *Acartia hudsonica* or *Acartia tonsa* females were isolated from the contents of the plankton tow and placed individually in petri dishes for egg production and hatching success assays. During these assays, females were fed maximum rations of mixture of *Tetraselmis spp*. and *Thalasiosira weissfloggi*. This diet has been used to maintain large cultures of these copepods for many generations in our lab (Dam *et al*., 2021; Sasaki & Dam, 2021). Females produced eggs for three days, after which the female was removed. Eggs were given three additional days to hatch. Egg production (the total number of eggs produced per female per day), the hatching success (percentage of eggs produced that hatched), and the total offspring production (the number of nauplii produced per female per day) were measured for each individual. These assays were performed across a range of temperatures (10-30°C for *A. tonsa* and 4-24°C for *A. hudsonica*) using stand up incubators (Fisher Scientific Model #3720) Thermal performance curves (TPCs) for egg production, hatching success, and production were modeled using a Gaussian equation (Padfield *et al*., 2021). TPC parameters (maximum trait values and thermal optima) were extracted from these model fits. All analyses were performed in R version 4.1.3.

**Table 1:** Details for the field thermal performance curve collections. The date and water temperature (°C) at the time of collection are provided for each group, along with which species was collected. The exact date for the June 2015 collection of *Acartia hudsonica* was not available.

| Month | Date | Temperature | Species |
| --- | --- | --- | --- |
| January | January 21st 2015 | 2.1 | *Acartia hudsonica* |
| February | February 21st 2015 | -0.4 | *Acartia hudsonica* |
| March | March 16 2015 | 4.0 | *Acartia hudsonica* |
| April | April 21st 2015 | 9.2 | *Acartia hudsonica* |
| May | May 14th 2015 | 12.0 | *Acartia hudsonica* |
| June | June 2015 | 18.6 | *Acartia hudsonica* |
| July | July 29th 2014 | 20.0 | *Acartia tonsa* |
| August | August 13th 2014 | 18.0 | *Acartia tonsa* |
| September | September 11th 2014 | 18.0 | *Acartia tonsa* |
| October | October 22nd 2014 | 16.0 | *Acartia tonsa* |
| November 1 | November 4th 2014 | 14.5 | *Acartia tonsa* |
| November 2 | November 19th 2014 | 11.0 | *Acartia tonsa* |

Thermal survivorship curves were also determined for each of these collections by exposing individual females to an acute 24 hour heat shock across a range of temperatures. Individual females were placed in a 2.5 ml microfuge tube with 0.2 um filtered seawater, and moved to 15-well drybaths (USA Scientific). Each drybath was set to a single temperature, ranging from 17-28°C for *A. hudsonica* and 16-36°C for *A. tonsa*. Survivorship was checked after 24 hours. Survivorship was modeled using a logistic regression of individual survivorship against stress temperature. These survivorship curves were then used to estimate thermal tolerance (as LD50, the temperature of 50% mortality).

To determine whether TPC parameter variation tracked seasonal temperature changes, we regressed the TPC parameters against the temperature at the time of collection. A significant, positive relationship between collection temperature and TPC parameters (esp. thermal optimum and thermal tolerance) would indicate that TPCs varied in relation to the seasonal temperature cycle.

### Simulated Heat Waves

To examine transgenerational effects of heatwaves, we collected *Acartia tonsa* from the same site again in 2015 for use in laboratory simulated heatwave experiments. In order to test the effects of a heatwave against the seasonally shifting baseline of ambient temperature, three collections were made before, during, and after peak environmental temperatures, corresponding to late June, late July, and early December (Table 2). These experiments examined both direct and transgenerational effects of heatwaves. To examine the direct effects, egg production rate, egg hatching success, and offspring production were measured for around 60 females per collection, split into two groups (control and heatwave). These assays were performed as described in the Field TPC section, with females isolated in individual petri dishes and provided with food *ad libitum*. The control group remained at a temperature near the current ambient temperature in Long Island Sound while the Heatwave group experienced temperatures 5°C above the ambient temperature (Table 2). During the simulated heatwave, traits were measured over two periods to examine the effects of short and long heatwave events -Days 1-3 and Days 5-7. Females were moved into petri dishes with fresh food solution on Days 3 and 5. Eggs produced during the intervening period were not examined.

**Table 2:**
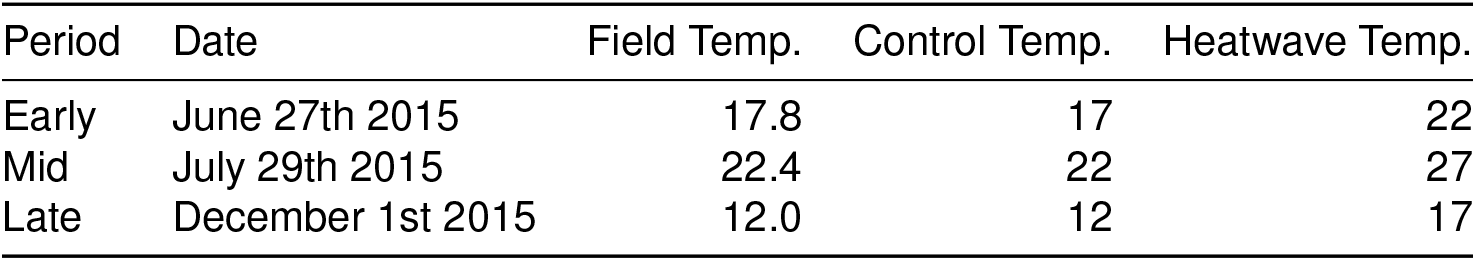
Details for the simulated heatwave collections. The date and water temperature at the time *A. tonsa* individuals were collected from the field is provided for each group, along with the temperatures used for the control and heatwave exposures.

| Period | Date | Field Temp. | Control Temp. | Heatwave Temp. |
| --- | --- | --- | --- | --- |
| Early | June 27th 2015 | 17.8 | 17 | 22 |
| Mid | July 29th 2015 | 22.4 | 22 | 27 |
| Late | December 1st 2015 | 12.0 | 12 | 17 |

We used effect sizes (Hedge’s g) to examine how heatwaves and the duration of exposure affected copepod performance (Gardner & Altman, 1986; Ho *et al*., 2019; Berner & Amrhein, 2022). Two sets of effect sizes were estimated within each collection: the effect of duration within treatments (long control vs. short control and long heatwave vs. short heatwave), and the effect of treatment (short heatwave vs. short control and long heatwave vs. long control). A negative effect size represents a reduction in the variable of interest as either 1) experimental duration increases, or 2) when copepods were exposed to heatwave conditions. Each effect size estimate also included a 95% Confidence Interval (CI), estimated by non-parametric bootstrapping (Ho *et al*., 2019). Hedge’s g was used instead of mean or median difference to account for the drastic seasonal differences in the magnitude and variation of egg production and hatching success observed between collections.

### Transgenerational Effects

In addition to the direct effects of heatwaves, we also examined the effects of parental exposure to increased temperature on offspring performance (i.e. -transgenerational effects). From the same collections described above, several hundred adult copepods were placed into each of eight 4L buckets of filtered seawater, which were split between the control and heatwave temperatures for that collection. Cultures were provided with food *ad libitum* and kept oxygenated using a small aquarium pump. Eggs were collected from each bucket following the same schedule as the direct effect experiments (on day 3 and day 7 for the short and long heatwave exposures, respectively). Eggs were discarded on day 5 to ensure all individuals reflected the correct exposure periods. For each collection, we therefore had four groups of offspring, those with parents exposed to control and heatwave temperatures for either 3 or 7 days.

These four groups of eggs were then further split into three groups which developed at either 12, 17, or 22°C. These rearing temperatures were used for all three collections, and reflect the temperatures observed at the times of collection. During development, individuals were maintained in 4L buckets of filtered seawater and fed *ad libitum*. After these individuals matured, body size and the three reproductive traits (egg production, hatching success, and production) were measured at the respective rearing temperature.

As for the F0 generation, we used effect size estimates (Hedge’s g) to examine how parental conditions affected offspring traits. For each seasonal collection we examined the effect of treatment within each developmental temperature (short heatwave vs. short control and long heatwave vs. long control). As offspring of the different batches of parents developed under identical conditions within each temperature, an effect of treatment in these comparisons indicates a transgenerational effect of parental exposure to heatwave conditions. As before, a negative effect size indicates a trait reduction when parents were exposed to heatwave conditions.

## Results

### Seasonal Variation in Field TPCs

TPCs for the two species were strongly diverged. There was also noticeable seasonal variation in the TPCs of *Acartia tonsa* (Table 3). *Acartia hudsonica* EPR TPCs had lower thermal optima and varied less across the season than those of *A. tonsa* (Figure 2). Across *A. tonsa* TPCs, EPR optimum temperatures tended to be higher in warmer months (Figure 2, Figure 3). Hatching success TPCs for *A. hudsonica* were also narrower and less seasonally variable than those of *A. tonsa*. Hatching success was higher for *A. tonsa* in warmer months than cooler months, regardless of incubation temperature. This is especially evident for the second November collection of *A. tonsa*, which exhibited very low hatching success at all but the highest temperatures used in the assays. When combined, the variation in EPR and HS TPCs yielded production curves that were highly variable in *A. tonsa*. Collections from warmer months generally had higher maximum production values and slightly higher optimum temperatures, although the narrow hatching success TPC of the second November collection produces a production TPC strongly skewed towards warmer temperatures, going against this trend.

**Table 3:**
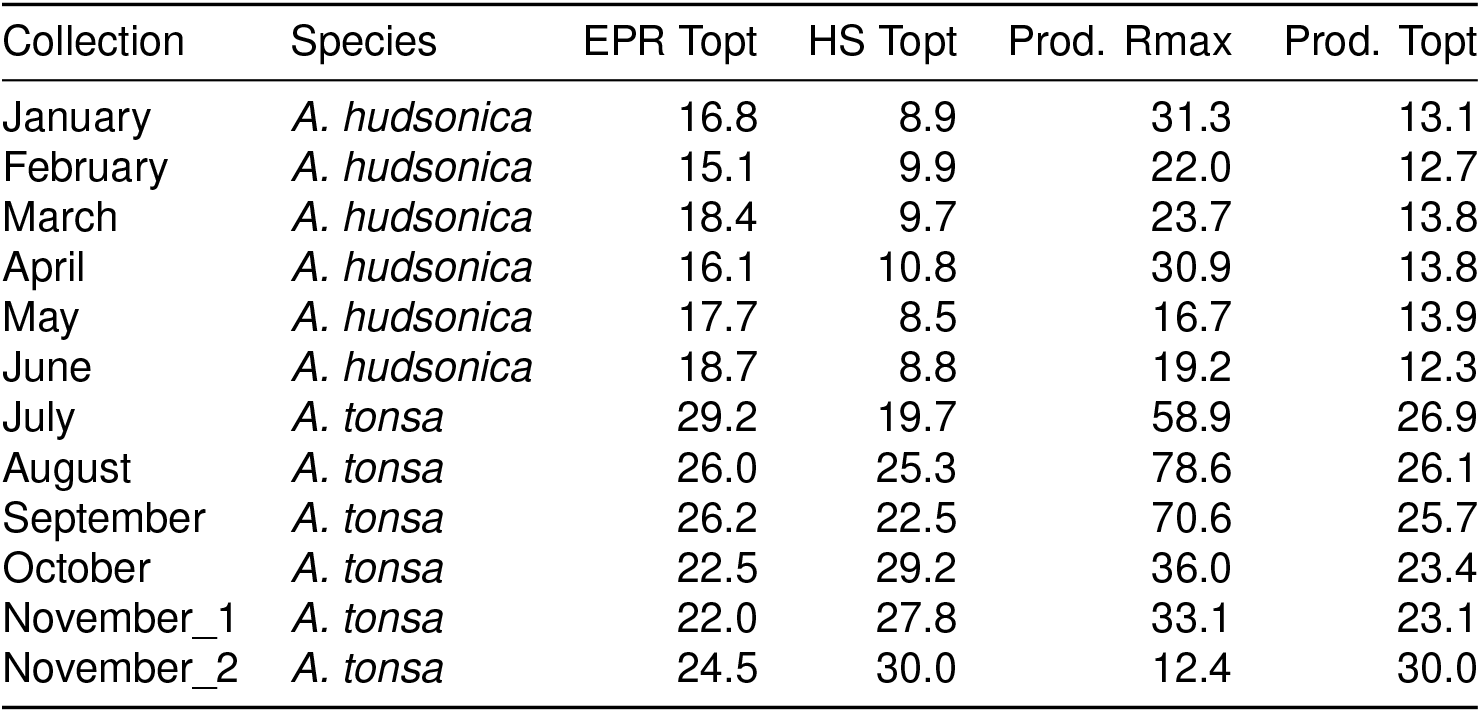
Subset of the thermal performance curve parameters for each collection (See supplemental file for full table, including 95% confidence intervals).

| Collection | Species | EPR Topt | HS Topt | Prod. Rmax | Prod. Topt |
| --- | --- | --- | --- | --- | --- |
| January | *A. hudsonica* | 16.8 | 8.9 | 31.3 | 13.1 |
| February | *A. hudsonica* | 15.1 | 9.9 | 22.0 | 12.7 |
| March | *A. hudsonica* | 18.4 | 9.7 | 23.7 | 13.8 |
| April | *A. hudsonica* | 16.1 | 10.8 | 30.9 | 13.8 |
| May | *A. hudsonica* | 17.7 | 8.5 | 16.7 | 13.9 |
| June | *A. hudsonica* | 18.7 | 8.8 | 19.2 | 12.3 |
| July | *A. tonsa* | 29.2 | 19.7 | 58.9 | 26.9 |
| August | *A. tonsa* | 26.0 | 25.3 | 78.6 | 26.1 |
| September | *A. tonsa* | 26.2 | 22.5 | 70.6 | 25.7 |
| October | *A. tonsa* | 22.5 | 29.2 | 36.0 | 23.4 |
| November_1 | *A. tonsa* | 22.0 | 27.8 | 33.1 | 23.1 |
| November_2 | *A. tonsa* | 24.5 | 30.0 | 12.4 | 30.0 |

**Figure 2:**
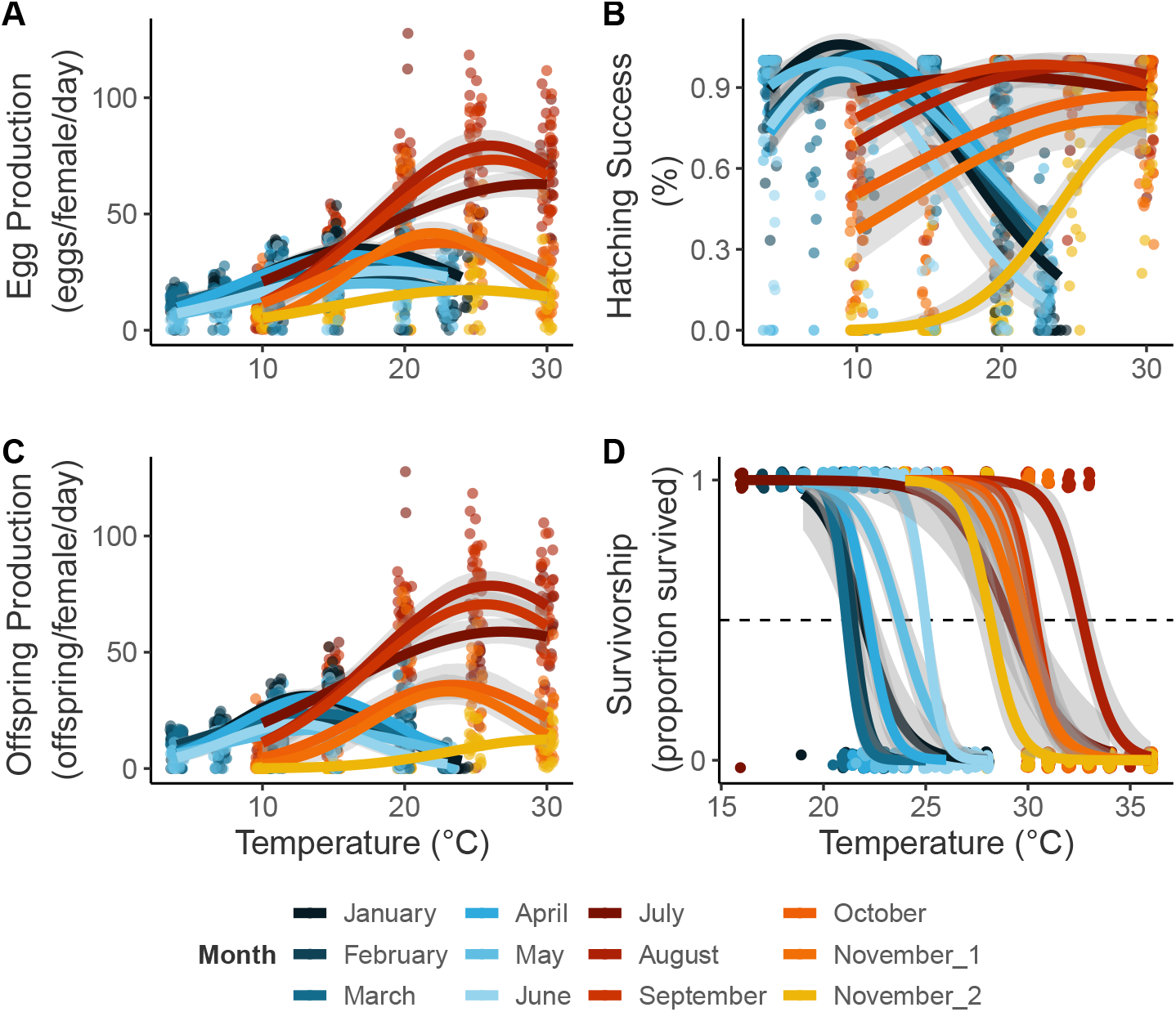
Thermal performance curves for each of the collections, each shown in a different color. Collections January through June examined Acartia hudsonica, while collections July through November examined A. tonsa. Panels A-C show points for each individual measurement along with the estimated Gaussian regression line. Panel D shows individual survivorship measurements following a 24-hour acute heat stress. Survivorship curves are estimated as logistic regressions.

**Figure 3:**
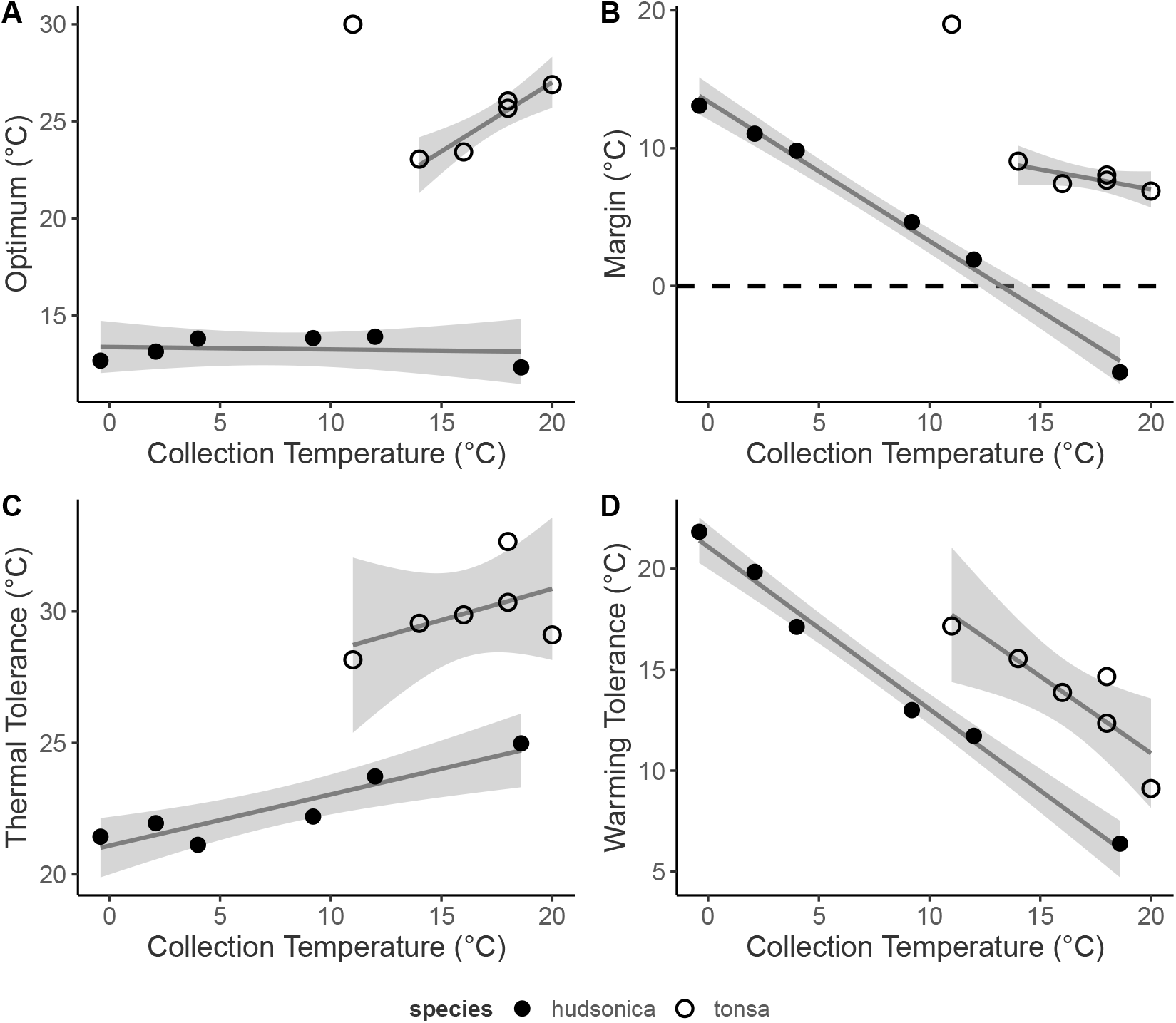
Relationships between (A) thermal optimum, estimated from the production thermal performance curves, and collection temperature; (B) the thermal safety margin (calculated as the difference between thermal optimum and collection temperature) and collection temperature; (C) thermal tolerance, measured as LD50, and collection temperature; and (D) warming tolerance (calculated as the difference between thermal tolerance and collection temperature) and collection temperature. The two Acartiid species are shown with different symbols. Linear regressions with confidence intervals are included.

Thermal optima of the offspring production TPCs were related to collection temperature for *A. tonsa* but not *A. hudsonica* (Figure 3). The values from the second November collection of *A. tonsa*, which had the highest production thermal optimum despite being collected at the lowest temperature, were excluded from this analysis. These copepods were collected at 11°C. This is around the threshold for resting egg production in *A. tonsa* (Holste & Peck, 2006). The extremely high estimated optimum temperature for production in this collection may therefore reflect the difference in hatching requirements between resting and subitaneous eggs. Thermal survivorship curves were variable in both species, with a range of LD50 values around 5°C in both species. Thermal tolerance increased with collection temperature in both species.

Vulnerability to heatwaves was estimated using two metrics: thermal safety margins for production (the difference between thermal optimum and collection temperature) and warming tolerance for survivorship (difference between thermal tolerance, LD50, and environmental temperature, Deutsch *et al*., 2008). Thermal safety margins of the two species responded differently to changes in collection temperature (Supp. Table 1). The invariant thermal optima for *A. hudsonica* production results in a strong decline in thermal safety margins as water temperatures increase (Figure 3; Supp. Fig. 1). Indeed, the warmest collection of *A. hudsonica*, occurring ∼19°C, appears to have been above the population’s thermal optimum. By contrast, the seasonally variable *A. tonsa* production TPCs resulted in relatively stable thermal safety margins (Figure 3; Supp. Fig. 1). Throughout its season of occurrence, *A. tonsa* maintained a safety margin of at least 5°C. In both species, warming tolerance responded in a similar manner to changes in collection temperature (Supp. Table 2; Supp. Fig. 1), with decreases in warming tolerance as water temperatures increased (Figure 3; Supp. Fig. 1). While collection temperatures never exceeded thermal tolerance for either species, the lower thermal tolerance of *A. hudsonica* translated to reduced warming tolerance relative to *A. tonsa*.

### Effects of Simulated Heatwaves

The second component of this project examined the effects of simulated heatwaves across generations in seasonal collections of *Acartia tonsa*. This began by assessing the impact of a simulated warming event on field collected (F0) individuals. We examined the effects of warming for 1-3 days (short duration) and 5-7 days (long duration) on EPR, HS, and offspring production (Figure 4; Supp. Fig. 2a). The impact of these warming events was assessed using an effect size estimate (Hedge’s g), comparing the warming to the control treatment. We focus here on the changes observed in body size and offspring production, which integrates the observed changes in egg production and hatching success.

**Figure 4:**
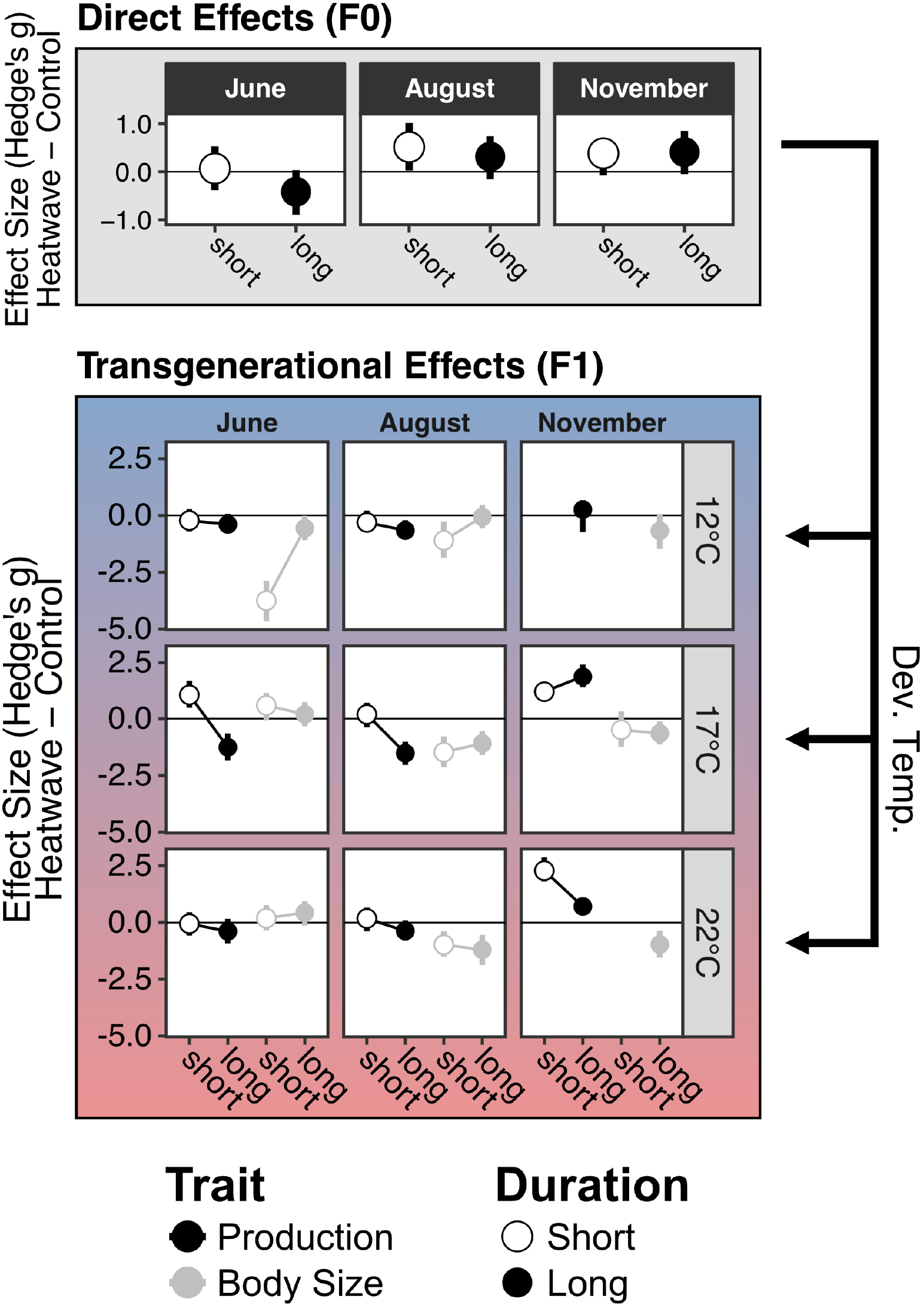
The effects of simulated heatwaves on production and body size. A) The direct effects of elevated temperature (field-caught F0 individuals), estimated as the effect size (Hedge’s g) calculated for the comparison between production by copepods exposed to heatwave and control treatments. A negative effect size indicates exposure to elevated temperature decreased production. The two different durations, short and long events, are shown with different symbols. B) The indirect, transgenerational effects of parental exposure to elevated temperature were also estimated as an effect size using Hedge’s g. Each panel shows data for one month and one developmental temperature. The different durations are again represented with different symbols, while offspring body size and production are shown in different colors. Here, a negative effect size indicates that parental exposure to elevated temperature decreased production or body size relative to individuals whos parents did not experience elevated temperatures.

Results were highly variable across collection and duration, but effect sizes tended to be fairly small (Hedge’s g between -1 and 1). For the June collection, there was a small increase in EPR in response to acute warming, and a small decrease in HS in response to the longer duration event. Production, integrating both EPR and HS, exhibited only a small decrease in the longer duration warming treatment. In the August copepods, there was a small increase in EPR and a small decrease in HS during both short and longer events. The increase in EPR was large enough, however, to result in a small increase in production during both short and long events. In November, warming resulted in a large increase in EPR regardless of the duration of the event (Hedge’s g > 1). There was no effect of warming on HS, however, limiting the increase in production.

We also examined the effect of warming duration (the comparison between long and short events within treatment) on the three traits. Here, we observed similar effects of duration on production in both control and warming groups, in all three months (Supp. Fig. 3); in all cases there were slight decreases in production over time for the June and August collections and an increase in production for the November collection. The similarities in the effect of duration between the control and heatwave treatments suggests that seasonal variation in the effects of warming may be more consequential than differences between short and long events (at least at the daily to weekly timescales examined here).

### Transgenerational Effects of Simulated Heatwaves

For the transgenerational experiments, comparing between control and heatwave treatments now examines the indirect effect of parental exposure to heatwaves on offspring performance. Results were again highly variable across developmental temperatures, parental exposure duration, and monthly collections. For the June and August collections, effect sizes were often similar across all three reproductive traits (Supp. Fig. 2b; see June copepods at 17°C and 22°C, and August copepods at 12 °C). When effect sizes differed between traits, EPR tended to be most strongly affected by parental exposure to warming (see June copepods at 18°C and August copepods at 17°C and 22°C). Differences in the effects of parental exposure to heatwaves on EPR and HS often ameliorated the overall effect size on offspring production, which was generally smaller than those for the other traits. There were, however, significant decreases in production driven by parental exposure to longer duration warming in offspring developed at 17°C for both June and August collections. A small increase in production resulted from parental exposure to short duration warming in the June copepods reared at 17°C. This was the only observation of a positive effect of parental exposure to warming on production for the June and August copepods.

Effects of parental exposure to elevated temperature on November copepods were drastically different from those observed in June and August copepods. Instead of primarily affecting EPR, parental exposure to heatwave conditions generally had large effects on HS in the November copepods (Supp. Fig. 2b). Additionally, unlike in June and August, there were no negative effects of parental exposure to warming on production values (Figure 3). However, nauplii in the 12°C developmental temperature group produced by parents exposed to short heatwaves did not successfully reach adulthood, potentially indicating strong lethal effects of parental exposure on offspring survival.

In all three collections, parental exposure to the heatwave conditions generally (but not always) reduced offspring body size. The effects of parental exposure to warming on production and body size were not correlated (Supp. Fig. 4). Offspring body size generally decreased with developmental temperature (Supp. Fig. 5). However, there were, at times, substantial differences in the observed temperature sensitivity of offspring body size between the two treatments; in June, parental exposure to heatwaves slightly decreased the temperature sensitivity of offspring body size, while a long heatwave during August increased the temperature sensitivity.

## Discussion

The rapid, intense warming associated with heatwaves and other extreme events may have dire consequences for ecological dynamics in aquatic systems. Responses to these acute events are determined by the position of thermal optima and limits relative to environmental temperatures. In taxa with short generation times, phenotypic plasticity and genetic differentiation may result in TPC variation that tracks ambient temperatures, maintaining a buffer against detrimental effects of heatwaves. Further, exposure to elevated temperatures during heatwaves may have indirect, transgenerational effects on offspring that are important to account for. We show that TPCs of *Acartia tonsa* but not *A. hudsonica* vary across seasons and that this variation allows the population of *A. tonsa* in Long Island Sound to maintain a relatively constant margin between optimum temperatures and ambient environmental temperatures. In *A. tonsa*, the mechanisms that produce seasonally variable TPCs, whether genetic or plastic, reduced the potential for heatwaves to adversely affect population dynamics and thus increase resilience in the face of climate change. We also used a series of simulated heatwave experiments to compare the transgenerational effects of short and long duration events throughout the year in *A. tonsa*. We observed significant variation in the transgenerational effects of heatwaves between collections, but results generally suggest only minimal deleterious effects of heatwaves on population dynamics across all seasonal collections. It may be that seasonal variation in thermal performance curves reduces parental stress during exposure to heatwave conditions, thus mitigating transgenerational effects on offspring performance.

Differing seasonal patterns in variation of thermal performance curves may affect which species will be most strongly affected by the increasing frequency and intensity of heatwaves. As a result of invariant TPCs, we observed strong seasonal variation in the vulnerability of *A. hudsonica* to heatwaves; as ambient temperature increases, it approaches the population’s relatively static thermal optimum. Indeed, late in the season of occurrence, ambient water temperatures actually exceed the thermal optimum. A heatwave at this point would be expected to strongly decrease population performance. By contrast, the variable TPC of *Acartia tonsa* produces relatively stable thermal safety margins, regardless of seasonal timing. As a result, we would predict that heatwaves would increase production by pushing environmental temperatures closer to the thermal optimum of this population. Importantly, the strong seasonal variation in TPCs is critical context for the widespread notion that *A. tonsa* constitutes a eurythermal species (Rahlff *et al*., 2017; González, 1974) -rather than a broad but fixed thermal performance curve, the large seasonal range of temperatures this population withstands is likely to be strongly affected by rapid shifts in the TPC. By focusing on short time scales this and other studies highlight that variation in the TPC is fundamental to shaping temporal occurrence in planktonic taxa (Sasaki & Dam, 2020; Anderson & Rynearson, 2020), and will likely play an important role in determining the response of taxa to climate change.

In the case of the two species examined here, the increasing incidence of heatwaves may reduce the seasonal occurrence of the cold-water dominant *Acartia hudsonica* as the thermal optimum is exceeded more frequently. *A. tonsa* performance, on the other hand, would be expected to increase during heatwaves due to the shift towards the thermal optimum. Latitudinal patterns in vulnerability to heatwaves might also be affected by the relative capacity for populations to adjust TPCs. At high latitudes, cold-specialists may lack the capacity to adjust TPCs (Peck *et al*., 2014), while at low latitudes, the presence of hard upper limits to physiological acclimation (Hoffmann & Sgrò, 2011; Stillman, 2003) may prevent shifts in TPCs to accommodate warming. In both cases, static TPCs would be expected to contribute to heightened vulnerability to heatwaves, similar to what we observed in *A. hudsonica*. An increased vulnerability to warming at low latitudes is observed at both the species- and population-levels (Barley *et al*., 2021; Sasaki *et al*., 2022; Nguyen *et al*., 2011; Morley *et al*., 2019). It is still an open question, however, how reduced capacity to adjust TPCs may affect the ability of tropical taxa to shift distributions polewards into more variable environments. While TPCs shifted towards higher temperatures will reduce vulnerability to extreme temperatures even during the warmest seasons, inflexible TPCs may result in significant fitness costs during cooler seasons due to a reduction in organismal performance. Additional work is needed on how the relative capacity for variation in thermal performance curves may shape broader biogeographic range shifts by modifying vulnerability to both high and low temperatures.

The transgenerational effects of heatwaves we observed varied in two main respects -the effect of the duration of parental exposure to increased temperatures across offspring developmental temperature treat-ments, and the pattern of this variation across collections. These results indicate that the effects of parental exposure to heatwaves are highly context specific, reinforcing that caution is warranted when extrapolating the results of both acute warming and short heatwave events from laboratory experiments to the response of natural populations. However, the general lack of strong detrimental effects does suggest that seasonal variation in TPCs may reduce transgenerational effects of heatwaves by preventing parental stress during these events.

Against the background variation in the effects of parental exposure to heatwaves across collections, however, there were at least superficial similarities between the effects in the June and August collections, and strong differences between these and the November copepods. Past work has shown that seasonal variation in thermal performance has a genetic basis in *A. tonsa* (Sasaki & Dam, 2020). This highlights the importance of considering not only seasonal variation in TPCs, but the mechanistic underpinnings of the variation. In scenarios where seasonal variation is entirely due to plasticity, we would expect broad similarities in the transgenerational effects of heatwaves regardless of offspring developmental temperature. However, where variation in TPCs is produced by genetic changes, transgenerational effects may also differ, reflecting the interplay between parental stress, developmental effects, and differing genetic backgrounds. This context dependence is a crucial yet largely unexplored dimension of the response of marine taxa to heatwaves.

The increasing frequency of heatwaves is just one of the aspects of climate change, and the ultimate impacts of heatwaves on population dynamics will represent the cumulative effects of multiple factors including, acidification, deoxygenation, and changes in food quality and quantity. These multi-stressor interactions have important consequences on evolutionary adaptation (Dam *et al*., 2021), and may play similar roles in the seasonal vulnerability to heatwaves. In particular, animals in our experiments were fed *ad libitum* and may have been able to maintain the energetic demands of a robust stress response via increased consumption. Alterations of thermal performance curves by reduced food availability may have strong impacts on population responses to increased temperatures (Huey & Kingsolver, 2019). We also suggest that future work examine how maternal effects may alter offspring responses to recurring heatwaves (Loveridge & Lucas, 2023; Minuti *et al*., 2022). In addition to the organismal basis for observed changes in thermal performance curves (plasticity vs. seasonal genetic differentiation for example), it will be important to identify the environmental drivers for patterns in variation over these relatively short timescales in order to better contextualize the nature of physiological stress (Dowd & Denny, 2020).

## Acknowledgements

We thank Zair Burris for her assistance with data collection. This study was funded by grants from Connecticut Sea Grant (R/LR-25) and the National Science Foundation (OCE-1947965) awarded to H.G.D., and a NSF postdoctoral fellowship (OCE-2205848) awarded to M.C.S.

## Supplementary Material

### Supplementary Tables

**Table S1:** ANOVA results for a linear regression examining the relationship between thermal safety margin and collection temperature for the two Acartiid species examined. The model included collection temperature and species as factors, along with their interaction.

|  | Sum Sq | Df | F value | Pr(>F) |
| --- | --- | --- | --- | --- |
| (Intercept) | 451.696 | 1 | 1017.230 | 0.000 |
| temperature | 256.400 | 1 | 577.418 | 0.000 |
| species | 0.023 | 1 | 0.052 | 0.826 |
| temperature:species | 10.034 | 1 | 22.596 | 0.002 |
| Residuals | 3.108 | 7 | NA | NA |

**Table S2:** ANOVA results for a linear regression examining the relationship between warming tolerance and collection temperature for the two Acartiid species examined. The model included collection temperature and species as factors, along with their interaction.

|  | Sum Sq | Df | F value | Pr(>F) |
| --- | --- | --- | --- | --- |
| (Intercept) | 1121.518 | 1 | 885.457 | 0.000 |
| temperature | 162.123 | 1 | 127.999 | 0.000 |
| species | 4.554 | 1 | 3.596 | 0.095 |
| temperature:species | 0.082 | 1 | 0.065 | 0.805 |
| Residuals | 10.133 | 8 | NA | NA |

### Supplementary Figures

**Figure S1:**
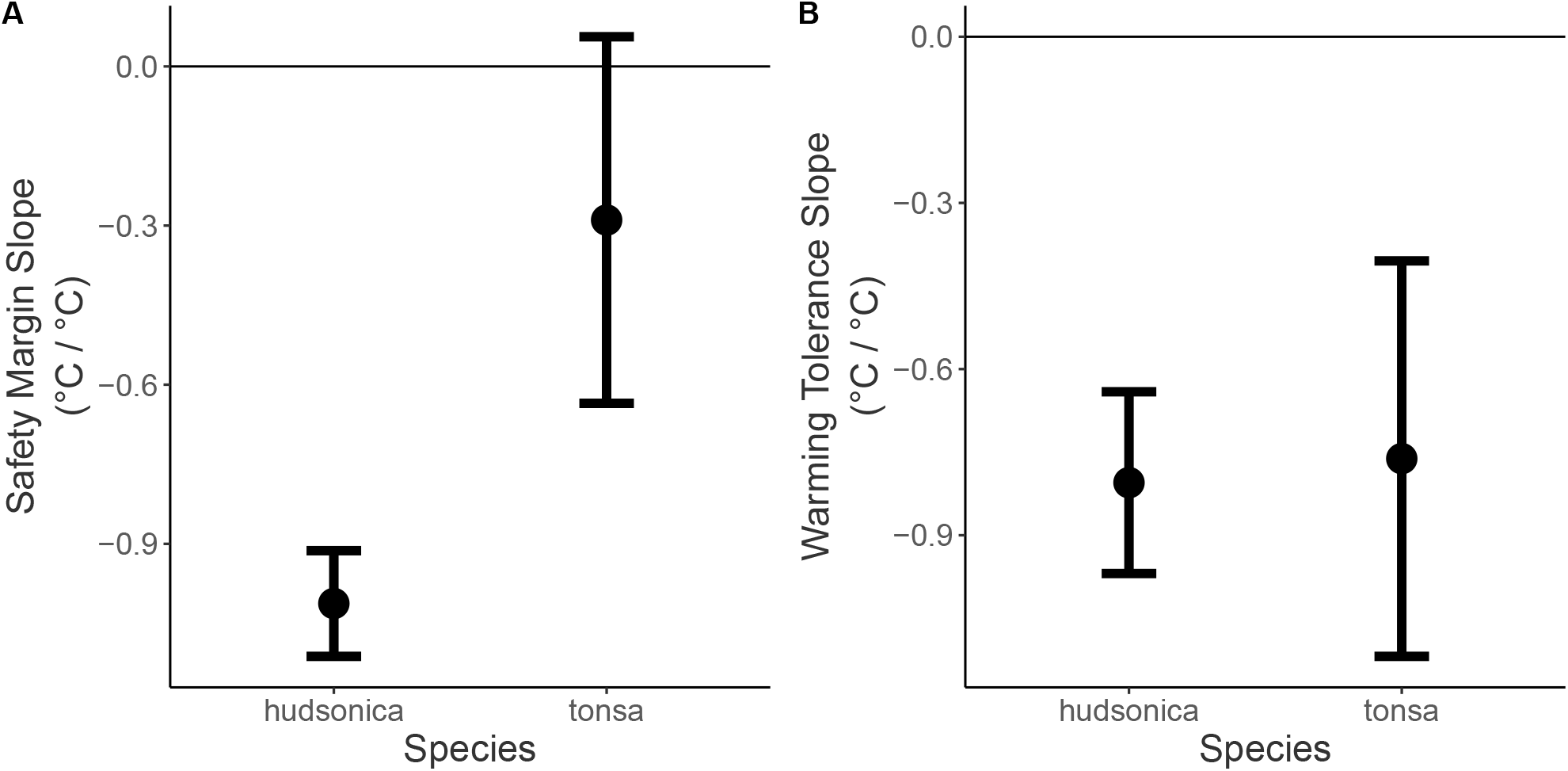
Comparison of the A) safety margin vs. collection temperature slopes and B) warming tolerance vs. collection temperature slopes for the two Acartia species, +/− 95 percent confidence intervals. Slopes represent the change in metric per °C change in collection temperature.

**Figure S2:**
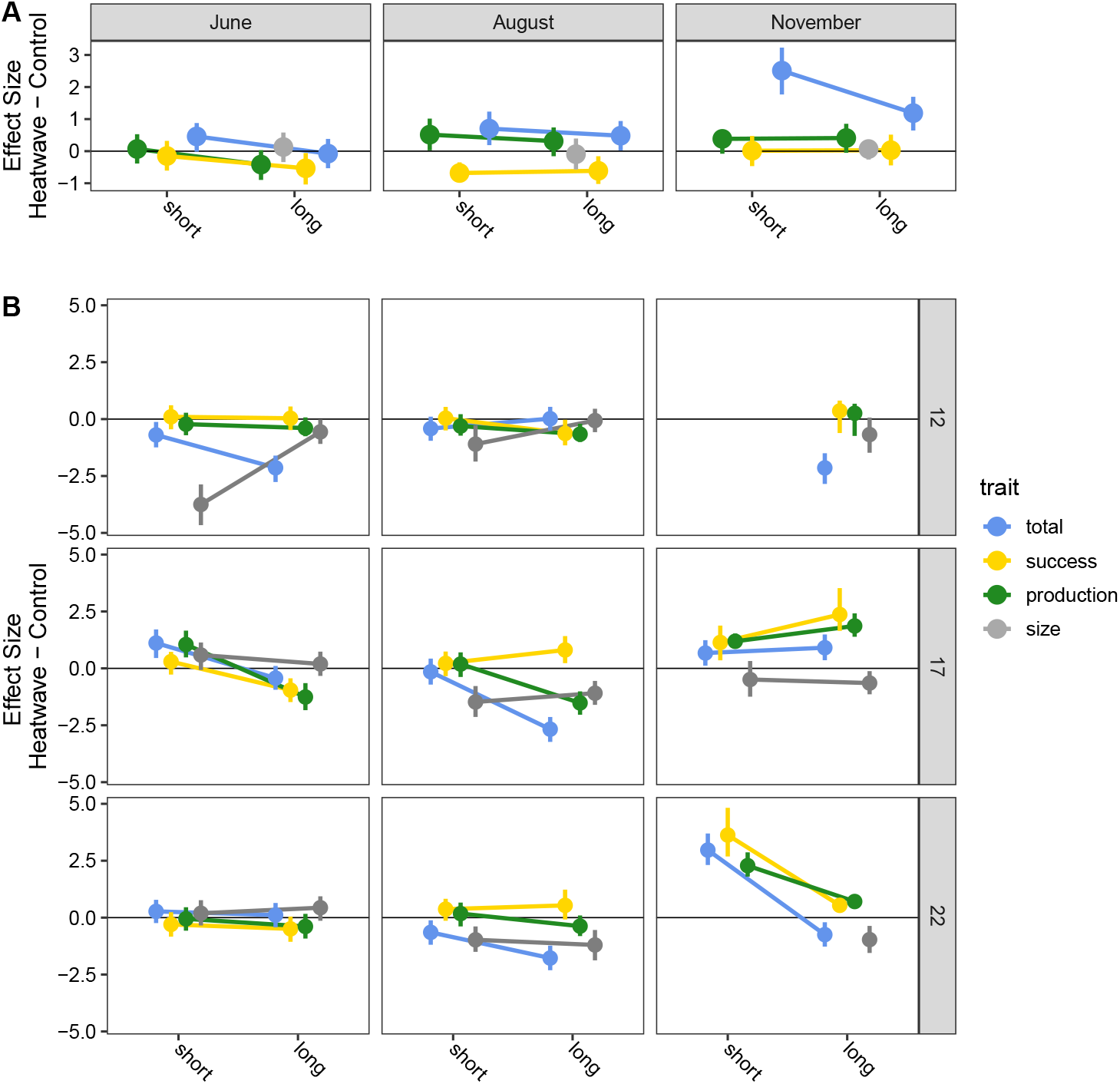
The effects of simulated heatwaves on all four measured traits: body size, production, hatching success, and egg production rate. A) The direct effects of elevated temperature (field-caught F0 individuals), estimated as the effect size (Hedge’s g) calculated for the trait comparisons between copepods exposed to heatwave and control treatments. A negative effect size indicates exposure to elevated temperature led to trait decreases. B) The indirect, transgenerational effects of parental exposure to elevated temperature were also estimated as an effect size using Hedge’s g. Each panel shows data for one month and one developmental temperature. Here, a negative effect size indicates that parental exposure to elevated temperature decreased production or body size relative to individuals whos parents did not experience elevated temperatures. In all cases, each of the four traits are shown in different colors.

**Figure S3:**
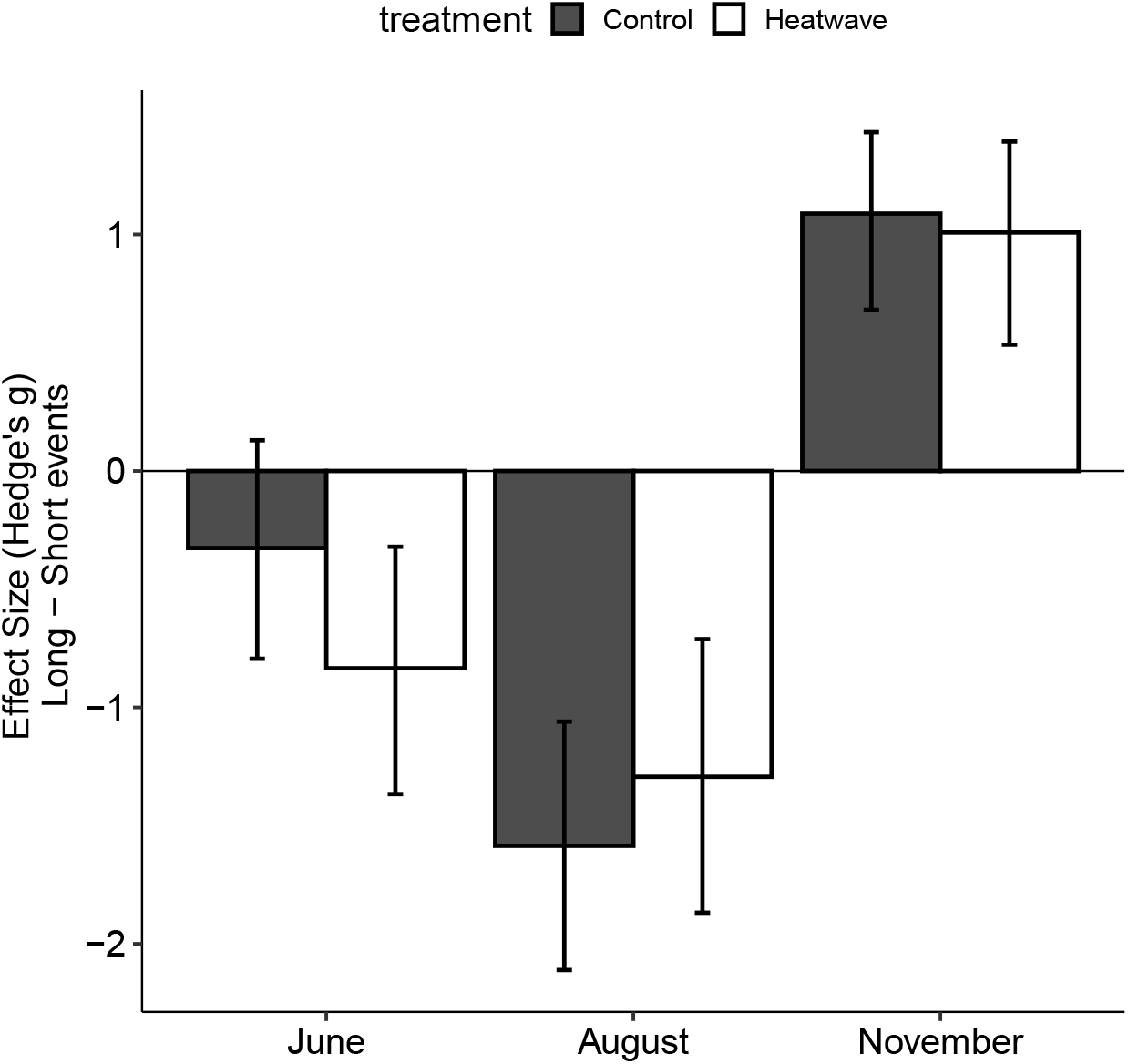
Effect size comparisons between short and long duration events for each collection, along with 95 percent confidence intervals. The different treatments, control and heatwave, are shown with different colors. A negative effect size indicates that the observed production value within a treatment decreased over time.

**Figure S4:**
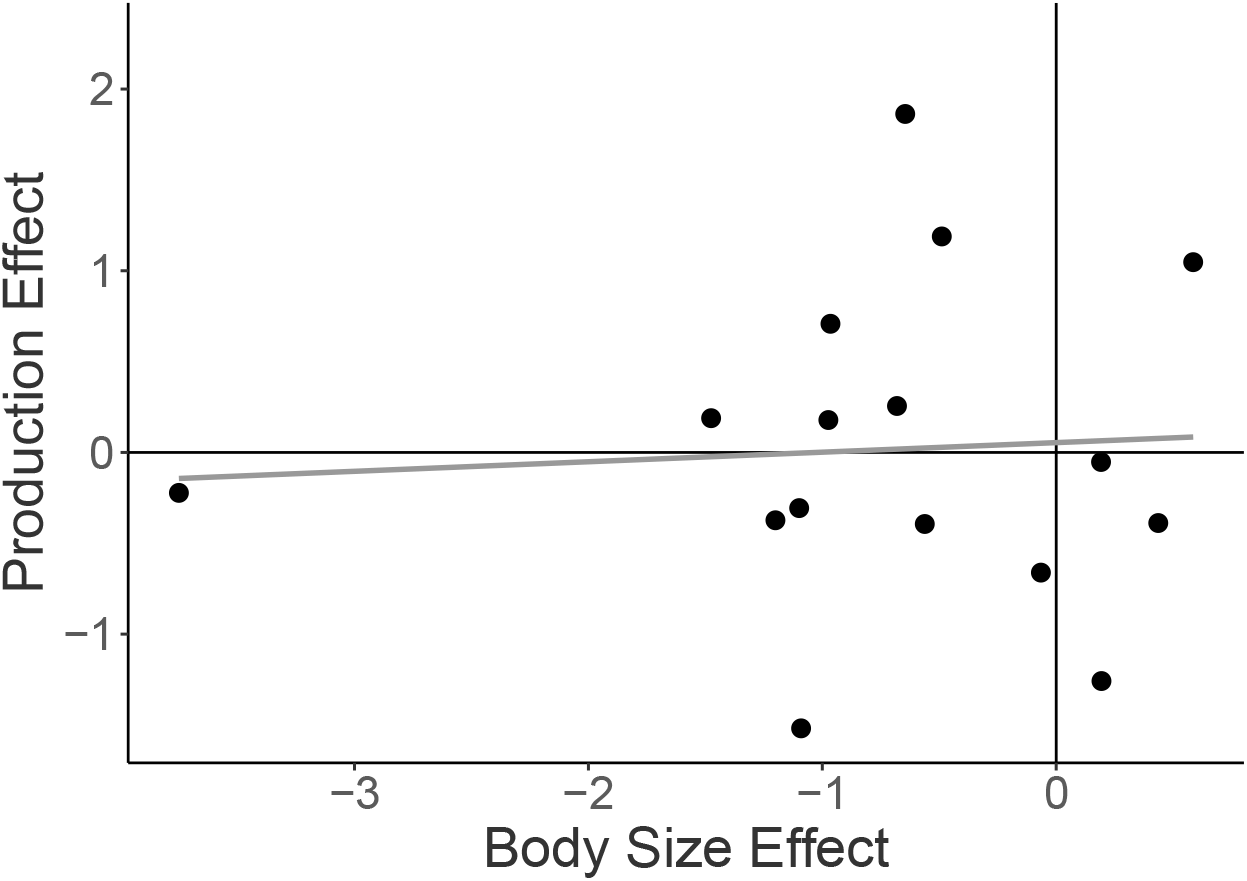
The relationship between the effects of parental exposure to warming on offspring body size and production. No significant relationship is observed.

**Figure S5:**
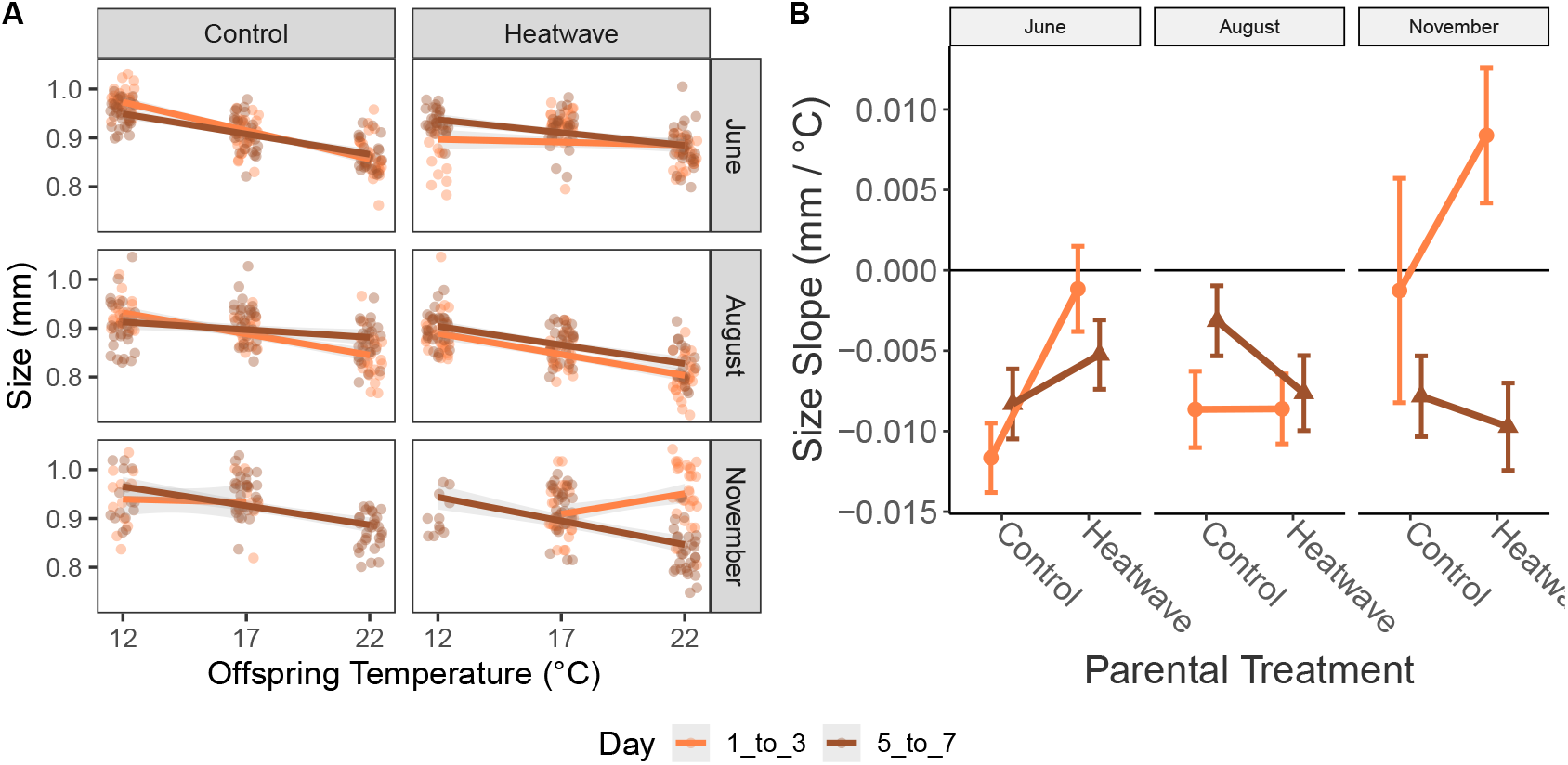
A) The relationship between offspring developmental temperature and adult body size. Each panel includes individual measurements for one treatment within a collection. Data for offspring of parents exposed to different duration events are shown in different colors. B) The slopes of body size vs. temperature shown in the panel A. A negative slope indicates that body size decreases with increasing temperature. The different durations are again shown with different colors.

