## Supplementary material for "Seasonally variable thermal performance curves prevent adverse effects of heatwaves": Table 3

| Collection | Species | EPR Max. Rate | EPR Optimum Temp. | HS Max. Rate | HS Optimum Temp. | Offspring Production Max. Rate | Offspring Production Optimum Temp. |
|---|---|---|---|---|---|---|---|
| January | *A. hudsonica* | 35.14 [31.9 - 38.6] | 16.77 [15.7 - 18.3] | 1.06 [1 - 1.1] | 8.87 [8.1 - 9.5] | 31.26 [27.9 - 34.1] | 13.15 [12.6 - 13.8] |
| February | *A. hudsonica* | 24.5 [21.5 - 27.7] | 15.13 [13.8 - 16.9] | 1.02 [1 - 1.1] | 9.95 [9.1 - 10.7] | 21.96 [19.4 - 25] | 12.68 [11.7 - 13.5] |
| March | *A. hudsonica* | 26.84 [23.5 - 29] | 18.44 [16.2 - 23] | 0.94 [0.9 - 1] | 9.67 [8 - 10.9] | 23.67 [20.7 - 26.2] | 13.81 [13 - 14.8] |
| April | *A. hudsonica* | 32.37 [29.9 - 34.8] | 16.1 [15.4 - 16.9] | 1.02 [1 - 1.1] | 10.8 [10 - 11.5] | 30.86 [28.4 - 33.5] | 13.84 [13.3 - 14.3] |
| May | *A. hudsonica* | 20.12 [18.4 - 21.9] | 17.66 [16.5 - 19.9] | 1 [0.9 - 1] | 8.51 [6.5 - 9.8] | 16.7 [14.6 - 18.9] | 13.91 [13 - 14.9] |
| June | *A. hudsonica* | 25.55 [22.4 - 28.2] | 18.66 [16.7 - 23] | 0.96 [0.9 - 1] | 8.81 [7.6 - 9.7] | 19.24 [15.9 - 22.5] | 12.34 [11.5 - 13.3] |
| July | *A. tonsa* | 62.85 [58.1 - 69.6] | 29.16 [25.8 - 30] | 0.94 [0.9 - 1] | 19.67 [10.3 - 27.7] | 58.86 [53.6 - 64.7] | 26.89 [24.5 - 30] |
| August | *A. tonsa* | 79.37 [71.5 - 85.3] | 25.97 [24.8 - 27.8] | 0.98 [0.9 - 1] | 25.3 [22.5 - 30] | 78.62 [71.6 - 85] | 26.05 [24.9 - 28.1] |
| September | *A. tonsa* | 73.34 [67.3 - 78.3] | 26.24 [25 - 28.2] | 0.99 [0.9 - 1] | 22.5 [20.9 - 25] | 70.62 [64.7 - 75.7] | 25.67 [24.6 - 27.1] |
| October | *A. tonsa* | 37.53 [32.1 - 42.5] | 22.53 [21.4 - 24] | 0.87 [0.7 - 1] | 29.2 [23.2 - 30] | 36.02 [28.7 - 43.5] | 23.43 [22.2 - 24.7] |
| November_1 | *A. tonsa* | 41.77 [36.8 - 46.7] | 21.98 [21.1 - 22.8] | 0.78 [0.7 - 0.9] | 27.76 [23.5 - 30] | 33.15 [27.6 - 40.3] | 23.06 [21.9 - 24.3] |
| November_2 | *A. tonsa* | 17.25 [14.9 - 19.8] | 24.47 [22.4 - 28.2] | 0.77 | 30 | 12.45 | 30 |
